# Complete genome assembly of the *Wolbachia* endosymbiont of the horn fly *Haematobia irritans irritans:* a supergroup A strain with multiple horizontally acquired cytoplasmic incompatibility genes

**DOI:** 10.1101/836908

**Authors:** Mukund Madhav, Rhys Parry, Jess A.T. Morgan, Peter James, Sassan Asgari

## Abstract

The horn fly, *Haematobia irritans irritans*, is a hematophagous parasite of livestock distributed throughout Europe, Africa, Asia, and the Americas. Welfare losses on livestock due to horn fly infestation are estimated to cost between USD 1-2.5 billion annually in North America and Brazil. The endosymbiotic bacterium *Wolbachia pipientis* is a maternally inherited manipulator of reproductive biology in arthropods and naturally infects laboratory colonies of horn flies from Kerrville, USA and Alberta, Canada, but has also been identified in wild-caught samples from Canada, USA, Mexico and Hungary. Reassembly of PacBio long-read and Illumina genomic DNA libraries from the Kerrville *H. i. irritans* genome project allowed for a complete and circularised 1.3 Mb *Wolbachia* genome (*w*Hae). Annotation of *w*Hae yielded 1249 coding genes, 34 tRNAs, three rRNAs, and five prophage regions. Comparative genomics and whole genome Bayesian evolutionary analysis of *w*Hae compared to published *Wolbachia* genomes suggests that *w*Hae is most closely related to and diverged from *Wolbachia* supergroup A strains known to infect *Drosophila* spp. Whole-genome synteny analyses between *w*Hae and closely related genomes indicates that *w*Hae has undergone convoluted genome rearrangements while maintaining high nucleotide identity. Comparative analysis of the cytoplasmic incompatibility (CI) genes of *w*Hae suggests two phylogenetically distinct CI loci and acquisition of another *CifB* homolog from phylogenetically distant supergroup A *Wolbachia* strains suggesting horizontal acquisition of these loci. The *w*Hae genome provides a resource for future examination of the impact *Wolbachia* may have in both biocontrol and potential insecticide resistance of horn flies.

**Importance:** Horn flies, *Haematobia irritans*, are obligate hematophagous parasites of cattle having significant effects on production and animal welfare. Control of horn flies mainly relies on the use of insecticides, but issues with resistance have increased interest in development of alternative means of control. *Wolbachia pipientis* is an endosymbiont bacterium known to have a range of effects on host reproduction such as induction of cytoplasmic incompatibility, feminization, male killing, and also impacts on vector transmission. These characteristics of *Wolbachia* have been exploited in biological control approaches for a range of insect pests. Here we report the assembly and annotation of the circular genome of the *Wolbachia* strain of the Kerrickville, USA horn fly (*w*Hae). Annotation of *w*Hae suggests its unique features including the horizontal acquisition of additional transcriptionally active cytoplasmic incompatibility loci. This study will provide the foundation for future *Wolbachia-*induced biological effect studies for control of horn flies.

## Introduction

Flies from the genus *Haematobia* (Diptera: Muscidae) are obligate hematophagous ectoparasites of pastured cattle. Two prominent members of this genus are the horn fly, *Haematobia irritans*, distributed throughout Europe, Africa, Asia, and the Americas (1) and the buffalo fly, *Haematobia irritans exigua*, which is widespread throughout Asia and Australia (2). Blood-feeding behaviour from *H. i. irritans* results in severe welfare issues and economic losses to cattle industries with annual estimates of up to $US ~1 billion in North America and $US ~2.5 billion in Brazil (3–5). In Australia, *H. i. exigua* is estimated to cost the domestic cattle industry $AUS 98.7 million annually and is currently restricted to the northern part of the country (6). Control of *Haematobia* flies primarily relies on the use of chemical insecticides; however, reports of insecticide resistance suggest that alternative intervention strategies are required (2, 7, 8).

*Wolbachia pipientis* is an obligate, endosymbiotic, Gram-negative α-proteobacteria estimated to infect between 40-70% of terrestrial arthropods (9, 10). *Wolbachia* infection in insects is known to selfishly alter host reproductive biology to transmit and persist in the next generation (11). One mechanism that drives transgenerational *Wolbachia* persistence is known as cytoplasmic incompatibility (CI) (12, 13). In CI, mating between *Wolbachia*-infected male and non-infected female (unidirectional CI) or female infected with a different *Wolbachia* strain (bidirectional CI) results in embryo death (13). The commonly accepted model for CI is “mod/resc”. Here, *mod* stands for modification of sperm by a toxin in the *Wolbachia*-infected male, and *resc* for a rescue of sperm by an antidote present in the egg (12, 14). Cellular studies have linked early embryonic death with defects in first zygotic mitosis, irregular chromosomal condensation post-fertilisation, and delayed histone deposition in the earlier interphase cell cycle (15–18). Two parallel studies recently identified the molecular mechanisms underpinning CI. Using a combined genomic and transcriptomic approach, LePage et al. (2017) identified two genes, *cifA* and *cifB*, in the prophage WO of *w*Mel *Wolbachia* strain mediating CI (19). Whereas Beckmann et al. (2017) demonstrated two genes *cidA* and cidB, *cifA* and *cifB* homologues, underpinned CI in the supergroup B *Wolbachia* strain *w*Pip (20). Further experimental examination of the CI loci suggested a “Two-by-One” model, whereby the *cifA* gene works as the rescue factor, and *cifA* and *cifB* together instigated CI (21).

In addition to CI, other phenotypes of reproductive manipulation have been reported for *Wolbachia* including male-killing, parthenogenesis, and feminisation (13). *Wolbachia* has also been demonstrated to confer protection against RNA virus infection in dipteran hosts (10, 11). Both CI and the ability of *Wolbachia* to restrict RNA viruses form the basis for the deployment of *Wolbachia*-infected *Aedes aegypti* mosquito for the control of dengue fever and other arboviruses worldwide (22, 23).

In previous studies, *Wolbachia* have been found to replicate in higher density in organophosphate resistant *Culex pipiens* mosquitoes than susceptible individuals resulting in deleterious fitness effects (24, 25). However, no such association between insecticide resistance and *Wolbachia* density was observed in *Ae. aegypti* mosquitoes suggesting that interactions between the host insecticide resistance and *Wolbachia* dynamics is both host and *Wolbachia* strain dependent (26).

While *H. irritans* are not currently known to be vectors of pathogenic viruses in livestock, there exists significant interest in exploiting the CI phenotype of *Wolbachia* as a form of sterile insect technique in *H. i. exigua* in Australia. A comprehensive screen of *H. i. exigua* samples from 12 locations in Australia and also Bali, Indonesia did not detect *Wolbachia* (27). By comparison *Wolbachia* has been previously identified in many wild-caught populations of *H. i. irritans* from Mexico (28), field-caught and laboratory colonies from the USA (29, 30), both field-collected and laboratory colonies from Alberta, Canada (27, 31), and also from field-collected samples in Hungary (32).

The genome of the *H. i. irritans* Kerrville reference strain maintained at the USDA-ARS Knipling-Bushland U. S. Livestock Insects Research Laboratory (Kerrville, TX) was recently assembled using Pacific Biosciences (PacBio) SMRT technology and Illumina chemistries (33). Initial analysis of deposited sequencing data indicated that a large portion of the reads in both libraries shared similarity to the *Wolbachia* endosymbiont of *Drosophila simulans w*Ri strain (33, 34). During *H. i. irritans* genome assembly, the *Wolbachia* contigs were removed (personal communication Felix Guerrero; USDA-lab, US). Due to the intracellular nature of *Wolbachia* and presence of multiple insertion sequences within *Wolbachia* genomes, assemblies using only short-read chemistries often result in highly fragmented assemblies (35). Combining PacBio long-read sequencing and Illumina technologies has resulted in the closed and completed *Wolbachia* genome (35, 36).

In this study, we assembled and annotated a high-quality, circularised genome of the *H. i. irritans Wolbachia* strain (*w*Hae) and explored its phylogenetic relationship with the described *Wolbachia* strains, and the possibility of induction of CI by this strain based on what is known about the genes responsible for CI.

## Results and Discussion

### *w*Hae genome assembly, annotation and genome features

To extract and assemble the genome of *Wolbachia* from *H. i. irritans*, the genomic data from the Kerrville reference genome project (33) was trimmed and mapped against published *Wolbachia* genomes (34, 35, 37, 38) using BWA-MEM under relaxed mapping criteria (39). Initially, ~10 million of ~404 million paired-end Illumina reads and 128,203 of 4,471,713 (2.86%) PacBio reads mapped to representative supergroup A *Wolbachia* genomes. These reads were then extracted, and *de novo* assembled using Unicycler resulting in a singular, circularised draft assembly (40). Raw Illumina fastq reads were then iteratively mapped against this draft genome and polished using pilon (41). The final number of reads that mapped to the assembled *Wolbachia* genome were 140,429 out of 4,471,713 (3.1%) from the PacBio library, corresponding to an average coverage of ~187x, and 10,285,275 out of 404,202,898 (2.54%) from the paired-end Illumina libraries, corresponding to an average coverage of ~1280x. The final *w*Hae genome is 1,352,354bp with a GC content of 35.3%, which is similar to other previously assembled supergroup A *Wolbachia* strains (Table 1). The polished *w*Hae genome was then annotated using the NCBI prokaryotic genome annotation pipeline (42) which predicted that *w*Hae encodes for 1,419 genes with 1,249 protein-coding genes and 129 pseudogenes, with 56 containing frameshifts, 93 incomplete, 12 with an internal stop, and 31 with multiple problems. The RNA gene repertoire of the *w*Hae genome was identified to encode 34 tRNAs, three rRNAs (5s, 16s, and 23s), and also non-coding RNA genes such as RNase P RNA component class A (RFAM: RF00010), signal recognition particle sRNA small type (RFAM: RF00169), 6S RNA (RFAM: RF00013) and transfer-messenger RNA (RFAM: RF01849).

**Table 1:**
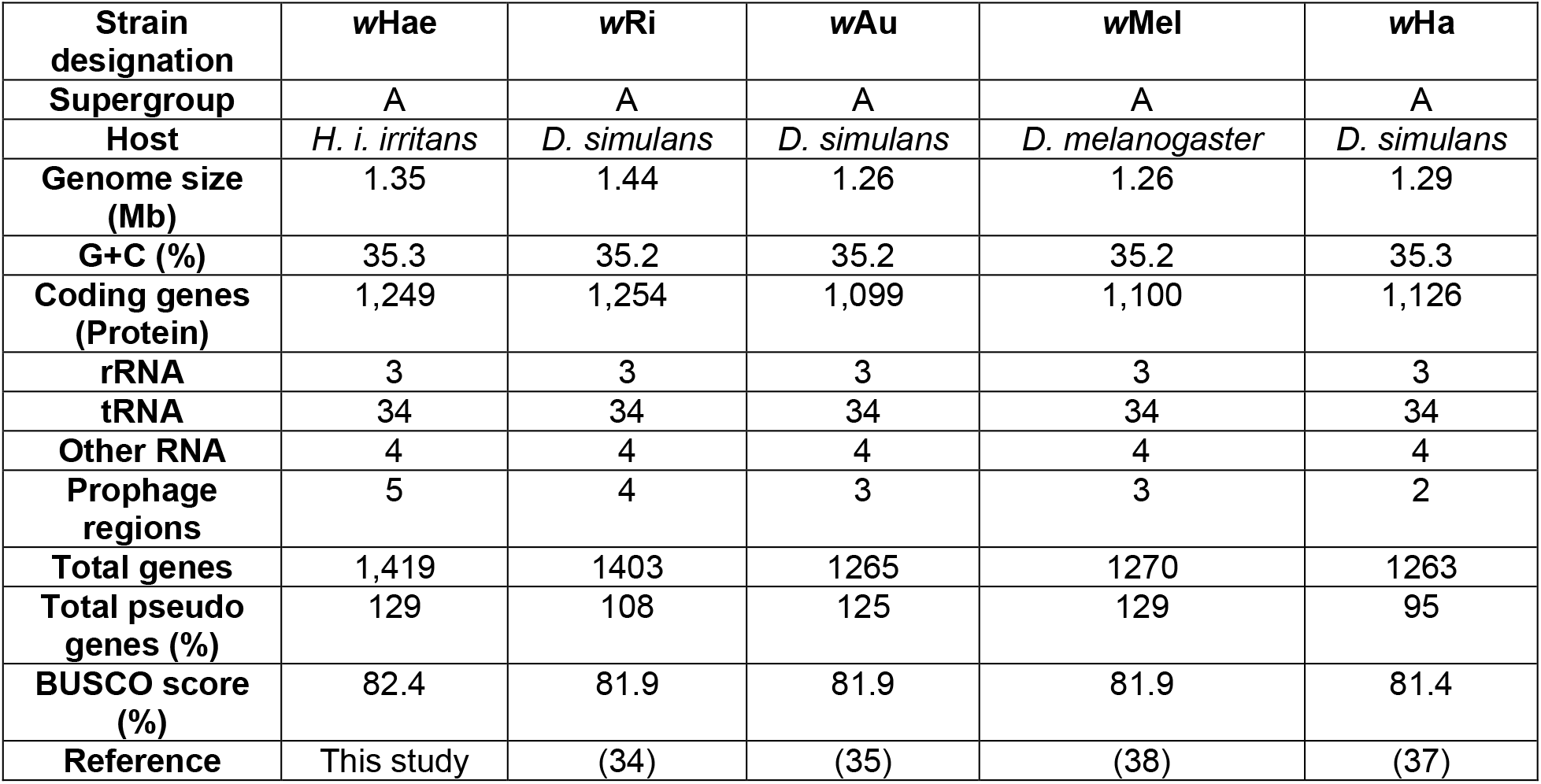
Genome features of complete supergroup A *Wolbachia* strains

| Strain designation | wHae | wRi | wAu | wMel | wHa |
| --- | --- | --- | --- | --- | --- |
| Supergroup | A | A | A | A | A |
| Host | *H. i. irritans* | *D. simulans* | *D. simulans* | *D. melanogaster* | *D. simulans* |
| Genome size (Mb) | 1.35 | 1.44 | 1.26 | 1.26 | 1.29 |
| G+C (%) | 35.3 | 35.2 | 35.2 | 35.2 | 35.3 |
| Coding genes (Protein) | 1,249 | 1,254 | 1,099 | 1,100 | 1,126 |
| rRNA | 3 | 3 | 3 | 3 | 3 |
| tRNA | 34 | 34 | 34 | 34 | 34 |
| Other RNA | 4 | 4 | 4 | 4 | 4 |
| Prophage regions | 5 | 4 | 3 | 3 | 2 |
| Total genes | 1,419 | 1403 | 1265 | 1270 | 1263 |
| Total pseudo genes (%) | 129 | 108 | 125 | 129 | 95 |
| BUSCO score (%) | 82.4 | 81.9 | 81.9 | 81.9 | 81.4 |
| Reference | This study | (34) | (35) | (38) | (37) |

Completeness of the *w*Hae genome was assessed by comparing the proteome against 221 single-copy orthologs derived from 1520 proteobacterial species in BUSCO pipeline (43). The BUSCO score for completeness of a model organism with a good reference genome is usually above 95%, but for the endosymbiotic bacteria with degenerated metabolic pathways BUSCO scores can vary between 50% to 95% based on the genome size, presence of repetitive elements in the genome and individual taxonomic placement (44). The completeness score for *w*Hae was 82.4%, which included 182 single-copy orthologs, two fragmented and 37 missing orthologs (Fig. S1), similar to five other completed *Wolbachia* genome projects (*w*Au, *w*Mel, *w*Ha, and *w*Ri).

Comparisons between the proteome of *w*Hae and four completed supergroup A *Wolbachia* strains (*w*Au, *w*Mel, *w*Ha, and *w*Ri) were carried out using the Orthovenn 2 web server (45). A total of 1136 orthologs were identified, of which all five strains shared 810 orthologs of which 782 single-copy genes were shared among all the strains with remaining specific to strains (Fig. S2). The *w*Hae genome has 1005 orthologs comprising of 1248 proteins, mostly involved in cell function and metabolism. Analysis of the proteome set of *w*Hae suggests that the “singleton” protein ortholog clusters exclusive to *w*Hae are transposable elements that are both present, but unannotated in the Genbank *Wolbachia* genome assemblies, or are exclusive to *w*Hae. These will be explored further below.

In addition to DNA sequencing data, we explored the transcriptional activity of *w*Hae in all life stages of *H. i. irritans* by mapping RNA-Seq data used to annotate the genome. As each sample was only sequenced once and poly-A enriched, it is difficult to make differential gene expression analyses with the data or infer *Wolbachia* tissue distributions. However, it appears *w*Hae is present and transcriptionally active in all life stages and all tissues dissected (Table 2). There is lower transcriptional activity in eggs and pupae than adults and also the highest normalised transcriptional activity was found in adult libraries at two hours post blood meal.

**Table 2:** Transcriptional activity of *w*Hae in all life stages of *H. i. irritans* Kerrville colony

| Run Accession | Sample | Reads Total | Reads mapped to wHae | Transcripts per million |
| --- | --- | --- | --- | --- |
| SRR6231662 | Malpighian tubules | 34,421,850 | 60,443 | 1,755.94 |
| SRR6231666 | Salivary gland | 27,461,942 | 2,580 | 93.94 |
| SRR6231664 | Adult 0h post feed | 44,820,832 | 181,392 | 4,047.04 |
| SRR6231671 | Adult 24h post feed | 47,070,858 | 188,823 | 4,011.46 |
| SRR6231665 | Adult 2h post feed | 51,829,048 | 827,413 | 15,964.27 |
| SRR6231670 | Adult 4h post feed | 50,512,062 | 205,766 | 4,073.60 |
| SRR6231654 | Egg 0h | 34,417,392 | 34,021 | 988.48 |
| SRR6231655 | Egg 2h | 38,158,656 | 48,477 | 1,270.40 |
| SRR6231660 | Egg 4h | 31,907,062 | 64,611 | 2,024.97 |
| SRR6231661 | Egg 9h | 33,832,868 | 61,261 | 1,810.69 |
| SRR6231658 | Midgut | 33,327,082 | 26,281 | 788.57 |
| SRR6231659 | Legs | 38,066,002 | 175,711 | 4,615.95 |
| SRR6231663 | Ovary | 39,372,642 | 51,187 | 1,300.06 |
| SRR6231668 | Pupae 1d | 53,975,828 | 274,698 | 5,089.27 |
| SRR6231669 | Pupae 3d | 51,859,710 | 377,067 | 7,270.90 |
| SRR6231667 | Testes | 87,918,073 | 1,167,195 | 13,275.93 |

### Phylogenetic placement of *w*Hae suggests close relationship between *Drosophila* spp. supergroup A *Wolbachia* strains

Since the discovery of *Wolbachia* within the gonads of the *Culex pipiens* mosquito, *Wolbachia* has taxonomically been considered a single species divided into 16 major supergroups (denoted A-Q) (46, 47). While the suitability of classifying the supergroups into a single *Wolbachia* species is the subject of ongoing debate (48, 49), a universal genotyping tool has been developed to demarcate supergroups based on multilocus sequence typing (MLST) of five ubiquitous genes (*gatB*, *coxA*, *hcpA*, *fbpA*, and *ftsZ*) (50). Although MLST clearly demarcates *Wolbachia* strains to supergroups, it fails to reliably discriminate strains within supergroups with high phylogenetic support. As such, a recent examination of these loci by Bleidorn and Gerth (2018) suggests that a number of alternative single copy loci outperform these five genes (50, 51). To construct a whole-genome phylogenetic analysis of *w*Hae, we used 79 of the 252 single copy orthologs from non-recombinant loci identified by Bleidorn and Gerth (2018) from 19 strains of *Wolbachia* (51). The phylogeny gives strong posterior probability support for the *Wolbachia w*Hae strain being basal to a clade containing *w*Rec, *w*Au and *w*Mel in supergroup A (Fig. 1).

**Figure 1:**
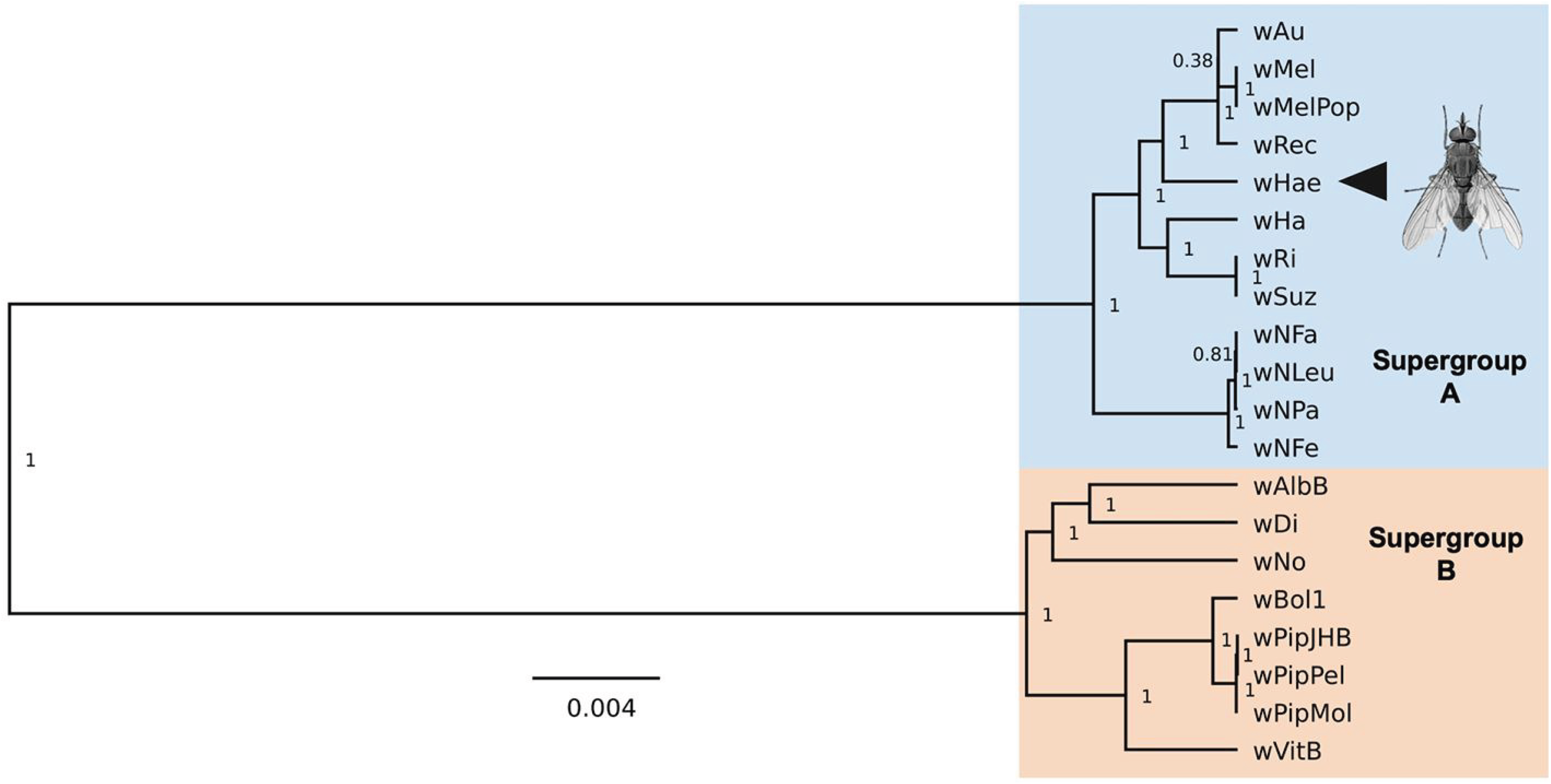
The *Wolbachia* endosymbiont of *Haematobia irritans irritans w*Hae is related to *Wolbachia* endosymbionts from *Drosophila* hosts. Maximum clade credibility (MCC) tree resulting from BEAST analyses of 79 concatenated recombination free gene loci of supergroup A and B *Wolbachia* strains previously identified by Bleidorn and Gerth (2018) resulting in an alignment of 49,68 bp. Posterior probability values are indicated at the nodes. *w*Hae indicated by an arrowhead and branch lengths represent the genetic distances.

Natural *Wolbachia* transfer between hosts can be cladogenic (*Wolbachia* acquired during the speciation of hosts), introgressive (transfer during mating between closely related host species), or horizontal (possibly via shared food and ecological niche, wounds and vectors) (52, 53). Concordance between the *Wolbachia* genome with the hosts mitochondrial and nuclear genome with consistent divergence time shows cladogenic transfer, whereas discordance suggests the possibility of horizontal transmission. Taxonomically, all Drosophilidae belong to the Ephydroidea superfamily of muscomorph flies, in which the *Wolbachia* strains *w*Au, *w*Ri, *w*Mel and *w*Rec have been identified. The *Haematobia* genus belongs to the subsection of Schizophora in the insect order Diptera, Calyptratae commonly referred to as the calyptrate muscoids (or simply calyptrates) (54). Evolutionary timescale analysis for the divergence of Ephydroidea and Calyptratae inferred from mitochondrial genes suggests that the most recent common ancestor of all *Haematobia* and *Drosophila* diverged sometime in the Palaeocene ~60 Million years ago (Mya) (55).

A number of phylodynamic analyses of *Wolbachia* genomes have attempted to reconstruct evolutionary timescales. However, there is limited concordance between analyses. Early analyses of the *ftsz* gene by Bandi et al. (1998) suggested that supergroups A-D diverged ~100 Mya (56), and a similar analysis was conducted by Werren et al. (1995) which suggested that the last common ancestor of supergroups A and B were approximately ~60 Mya (57). However, Gerth and Bleidorn (2016) proposed a much older divergence time between *Wolbachia* supergroups A and B of ∼200Mya (58). The Bayesian time to most recent common ancestor (TMRCA) analysis conducted by Gerth and Bleidorn (2016) on the clade encompassing all *Drosophila Wolbachia* strains was dated at 48.38 Mya, however, with a range of 110 – 16 Mya. Considering the mitochondrial divergence of *Haematobia* from *Drosophila* and early divergence of *Wolbachia* from supergroup A members (*w*Mel, *w*Ri, and *w*Rec) infecting *Drosophila* species, which are closely related to *w*Hae, due to the various timescale estimates and large range within the TMRCA we cannot rule out that the relationship between *w*Hae and other *Wolbachia* may be the result of codivergence. However, it is also possible that *Wolbachia* has been horizontally acquired in *H. i. irritans*.

### The *w*Hae genome has undergone convoluted genome rearrangements compared to other *Wolbachia* genomes

In bacterial genome evolution, horizontal gene transfer (59, 60) and genetic vehicles such as bacteriophages, plasmids or transposons (mobile element) (60–62) contribute to changes in the bacterial genome. Due to the intracellular niche of the endosymbiont, the evolution of *Wolbachia* genomes is highly dependent on bacteriophages, and transposable elements, with both contributing to sometimes as much as 21% of the genome (60).

Whole-genome comparisons of nucleotide synteny between *w*Hae and *w*Mel and *w*Ri were carried out using MAFFT v.7 (63). We did not analyse the synteny between *w*Rec (64) and wHae because the genome is fragmented and yet to be circularised. It appears that while *w*Hae maintains between 90-99% nucleotide identity with the other two strains, *w*Hae has undergone a high degree of genome rearrangement (Fig. 2A and B). In comparison, *w*Mel and *w*Ri show very similar genome arrangements (Fig. 2C). Similar genomic rearrangement has been previously seen while comparing *w*Pip and *w*Mel, *w*Mel and *w*Bm, *w*Uni and *wVitA* (65–67).

**Figure 2:**
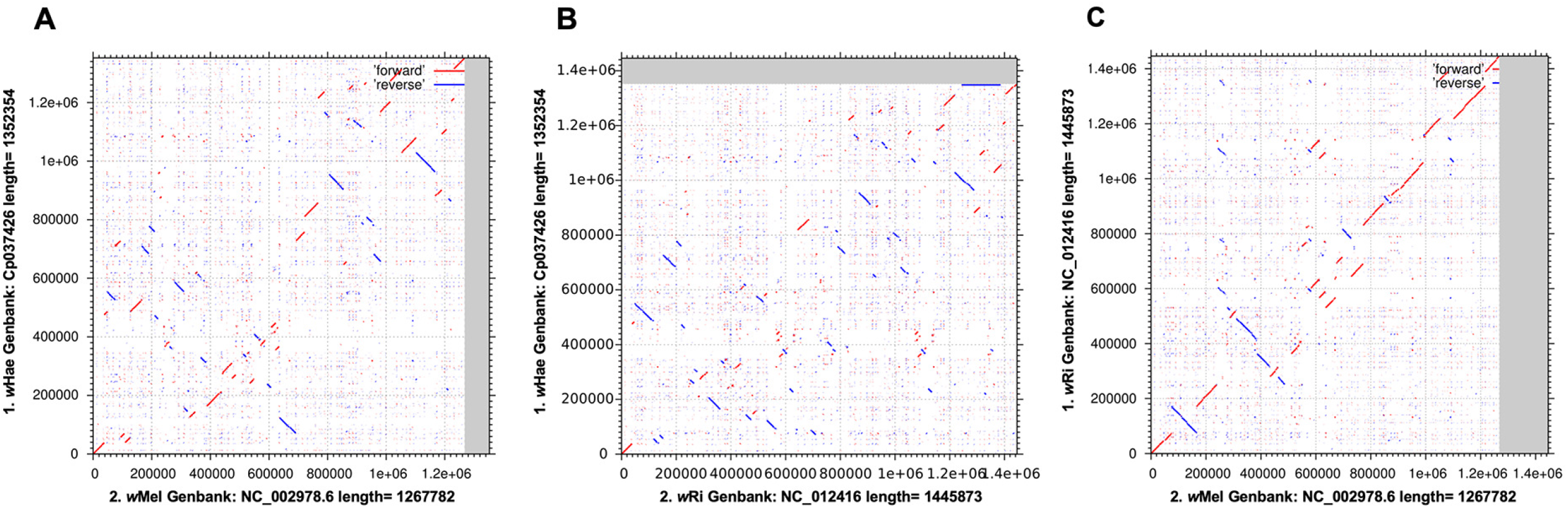
The *w*Hae genome has undergone convoluted genome rearrangements compared to other *Wolbachia* genomes. Genomes were compared using the MAFFT (v7) algorithm. Dot plots of LAST comparisons under (threshold score = 39 E=8.4e-11 A) *w*Hae genome compared to *w*Mel (Genbank: NC_002978.6) B) *w*Hae compared to *w*Ri (Genbank: NC_012416), and C) *w*Ri compared to *w*Mel. Similarities in the forward orientation (red) and similarities suggesting inversions (blue).

### Expansion of insertion sequence elements in *w*Hae genome is associated with a divergent CifB homologue

Insertion sequences (IS) are diverse transposable elements in bacterial genomes (60, 68). Considerable variation in the IS element composition in *Wolbachia* genomes is speculated to contribute to diversification or speciation of closely related strains, and IS elements can cause the disruption of protein coding genes leading to pseudogenes (35, 37). To compare the IS element load between *w*Hae and other supergroup A *Wolbachia*, *w*Ri, *w*Au, *w*Mel, *w*Ha, IS elements were identified and searched against the IS finder database using the ISsaga web server (68) (Supplementary File 1). A total of 283 ORFs related to IS elements were identified in the *w*Hae genome, including 61 complete ORFs and 150 partial IS elements. Maximum copies of IS elements were from IS630 (111 copies), which belong to the Tc1/mariner (Class II) transposon family, and ssgr IS1031 (109 copies), which is from the IS5 family. Comparative analyses between *w*Hae and other supergroup A *Wolbachia* strains identified 12 conserved IS families between all genomes IS66_ssgr_ISBst12, ISL3, IS5_ssgr_IS1031, IS4_ssgr_IS4, IS4_ssgr_IS231, IS3_ssgr_IS3, IS110, IS110_ssgr_IS1111, IS4_ssgr_IS50, IS630, IS481 and IS5_ssgr_IS903. However, two IS families were identified as exclusive to *w*Hae: IS5_ssgr_IS427, which has one complete ORF and three partial ORFs, and the IS5_ssgr_ISL2, with two partial ORFs. We manually extracted the IS5_ssgr_IS427 annotations and interestingly within one of the identified loci between positions 632,890 and 630,128, as annotated by the NCBI prokaryotic annotation pipeline as E0495_03250, a disrupted IS5-like element was found with the most closely related hit, based on BLASTn similarity (Query length:100%, Nucleotide identity: 80.39% E-value: 0), being the *Wolbachia* endosymbiont of *Brugia malayi* isolate TRS (Genbank ID: CP034333.1) (66). Immediately after this transposable fragment is the protein E0495_03245 (Fig. 3A), which BLASTp analysis of this 546aa protein appears to be a truncated CI factor CifB belonging to the *w*Ha *Wolbachia* endosymbiont of *Drosophila simulans* (Genbank ID: WP_144054595.1, Query cover: 98% Percentage similarity: 65.71% E-value: 0.0). We examined the transcriptional activity of this *cifB* gene by mapping the RNA-Seq data of all life stages to this region of the genome. Since only one paired reads mapped to this gene, it appears that the gene is transcriptionally silent (Fig. 3B). The length of IS elements varied between 174 to 1743 bp having a median size of 348bp. The total burden of IS elements on the *w*Hae genome is 115,692 bp, which is 8.55%. This is similar to the IS element percentage found in *w*Ri (9%) which is double that of the IS element load of *w*Mel (4.3%), *w*Ha (4.4%), and *w*Au (4.4%). This lineage-specific attainment and loss of IS elements, as well as length of the IS element, size and family distribution is well documented across *Wolbachia* strains (37). The association between IS elements conserved between *w*Hae from supergroup A, and *Wolbachia* from supergroup D and from the filarial nematode *Brugia malayi* is of particular interest. *B. malayi* is a filarial nematode that relies on a hematophagous mosquito host as a vector. Potentially, the gain of this IS element may have arisen through co-infection of *H. i. irritans* with a distantly related nematode species as it seems unlikely to have been independently lost in all other supergroup A genomes. While *H. i. irritans* is known to vector *Stephanofilaria* sp. nematodes (69), presence or absence of *Wolbachia* within these nematodes is yet to be characterised, and therefore formal testing of IS acquisition cannot be undertaken. Further assembly and genetic characterisation of filarial nematodes and their *Wolbachia* endosymbionts would allow for a better understanding of interaction between the *H. i. irritans*, *Stepahnofilaria* sp. and *Wolbachia*.

**Figure 3:**
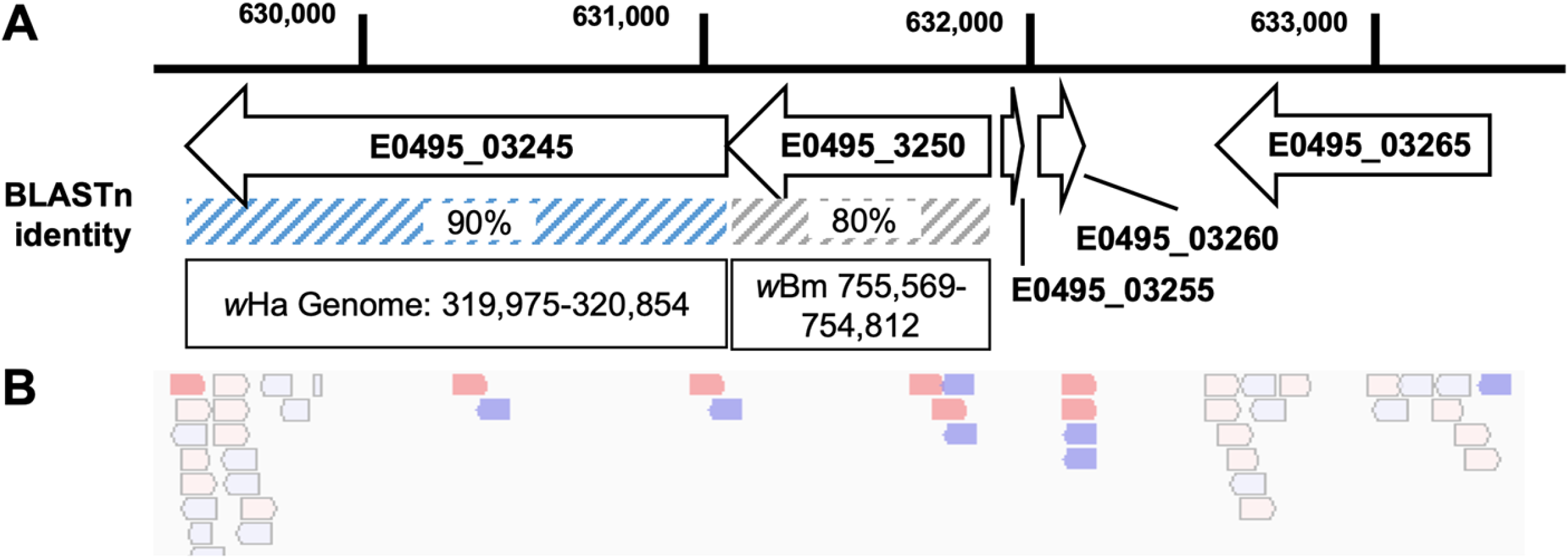
Expansion of IS elements in *w*Hae genome is associated with a transcriptionally silent *cifB* homologue. A) Schematic diagram of genomic loci in *w*Hae associated with the IS5_ssgr_IS427 IS family identified by ISsaga and BLASTn hits against the *w*Ha genome (Genbank ID:) and the *w*Bm genome (Genbank ID: CP034333.1). B) Transcriptional activity of the putative CifB homologue E0495_03245 was explored through pooling RNA-Seq reads originating from all tissues and developmental stages of all *H. i. irritans* libraries that were mapped to the *w*Hae genome. The resultant BAM files were visualized with Integrated Genomics Viewer (IGV v 2.5.2). Forward mapped reads are shown in red, reverse orientation reads are shown in blue. Light blue and red regions indicate a mapping quality number of 0 (MQ0) which indicates that the read maps to multiple regions on the genome.

### Comparative genomic analysis of prophage regions of *w*Hae

*Wolbachia* bacteriophages or prophages (WO) have been widely reported in strains from supergroup A, B and F, however, they have been lost in supergroup C and D strains (70). The tripartite relationship between *Wolbachia*–WO and arthropod hosts is of great interest as it has been shown that many genes located within prophage regions of *Wolbachia* genomes contain eukaryotic association genes and toxin-antitoxin modules (71), and also there is interest in utilising WO as a candidate for *Wolbachia* genetic transformation (70, 72). Using the Phaster web server, we identified five potential WO regions in the *w*Hae genome. The largest of which is a 60.8kb region designated as “intact” by Phaster with 68 ORFs from 359,527-420,415 having head, baseplate, tail, virulence genes and IS630 family transposons (73). The other four were ~7Kb incomplete prophage regions containing 10, 9, 12 and 8 ORFs positioned at 613,245-620,397, 859,203-866,672, 903,423-910,665, and 1,241,523-1,247,571 respectively in the *w*Hae genome. Supergroup A members *w*Mel, *w*Ri, *w*Au, and *w*Ha have between two to four variable WO phage regions with at least one presumed intact and other WO-like degenerated phage regions (34, 35, 37, 38). We compared the “intact” putative prophage region of WOHae with the predicted WO phage regions from *w*Mel (WOMelB) and completely sequenced WO phage region from *w*VitA (WOVitA), to identify the conserved region using reciprocal BLASTn analysis (38, 60, 74). The conserved phage regions were visualised using Easyfig (Fig. 4). We found that truncation of genes, insertion and deletion, or rearrangement of the genome has shifted the position of the base plate, tail and head region of WOHae. Ankyrin repeat domains (ANK) are involved in regulation of cell cycle, promotion of protein-protein interactions, and *Wolbachia-*induced reproductive phenotypes (75–77) and vary widely between strains (78). Total of four ANK were present in WOHae and WOMelB, but eight in WOVitA, suggesting a loss of ANK genes in different WO strains (77). WOHae had three and seven distinct hypothetical proteins of size 77-630aa compared to WOMelB and WOVitA, respectively. It has been suggested that these distinct hypothetical proteins could encode for the unique genes resulting in diversity amongst *Wolbachia* (77).

**Figure 4:**
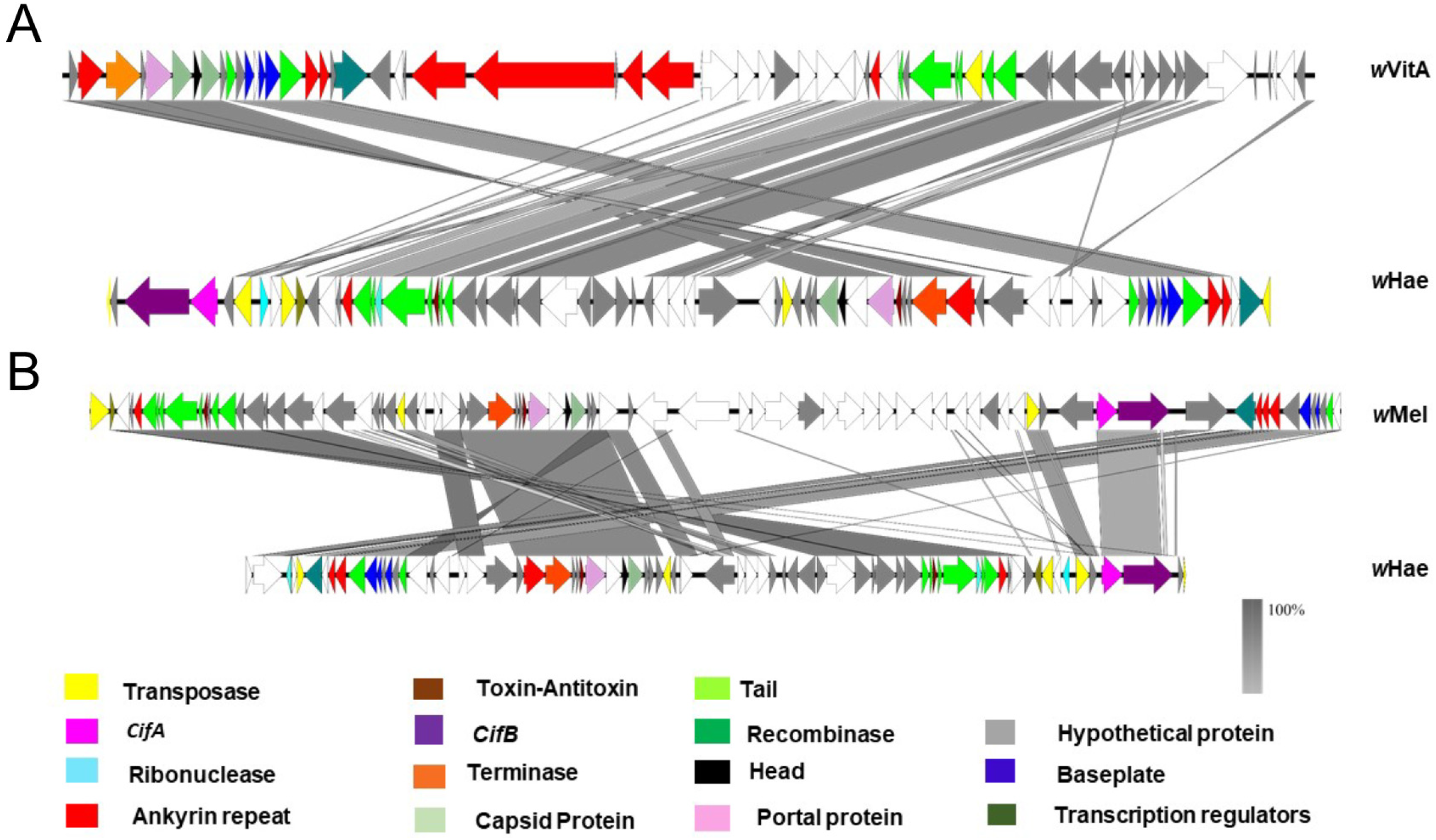
Gene order comparisons between WO prophages. Reciprocal BLASTn analyses of (A) Comparisons between WOVitA and WOHae, and (B) Comparisons between WOMelB and WOHae. Genomic loci in WO prophages were subjected to Easyfig and matching loci with max *E*-value (0.001) are indicated by grey shading. Annotations of genes are coloured based on automated NCBI annotation and manual PFAM protein database curation.

### Horizontal acquisition of *Wolbachia* cytoplasmic incompatibility loci in *w*Hae

To explore the genetic diversity of CI genes in *w*Hae, we explored orthologous clusters for the previously described CI genes. In addition to the truncated *cifB* (E0495_03245) gene, we found two complete and genetically distant CI loci in *w*Hae. With one located within the WOHae region (Gene ID: E0495_02160, E0495_02165) and the second CI locus (Gene ID: E0495_02270, E0495_02275) downstream of the WOHae region. BLASTp analysis of the predicted protein sequences (Table 3) indicated that these CI genes did not appear to be a duplication as previously reported for *w*Ri (34).

**Table 3:** Cytoplasmic incompatibility (CI) genes identified in *w*Hae and related protein

| Gene | Size | Position | Top Blastp hit, Query cover, Percentage identity, GenbankID |
| --- | --- | --- | --- |
| E0495_02160<br>cidA | 483aa | 414,360- 415,811 | cidA wPip, 100%, 82.89% AGR50404.1 |
| E0495_02165<br>cifB | 1132aa | 415,858- 419,216 | cifB wHa, 99%, 91.81%, WP_144054595.1 |
| cifA<br>E0495_02270 | 474aa | 441,815- 443,239 | cifA wMel 100%, 99.79%, WP_044471237.1 |
| cifB<br>E0495_02275 | 1166aa | 443,315 – 446,815 | cifA wMel, 99%, 99.40%, AYE93038.1 |
| E0495_03245 | 546aa | 629,487 – 631,127 | cifB wHa, 98% 65.71%WP_144054595.1 |

**Figure 5:**
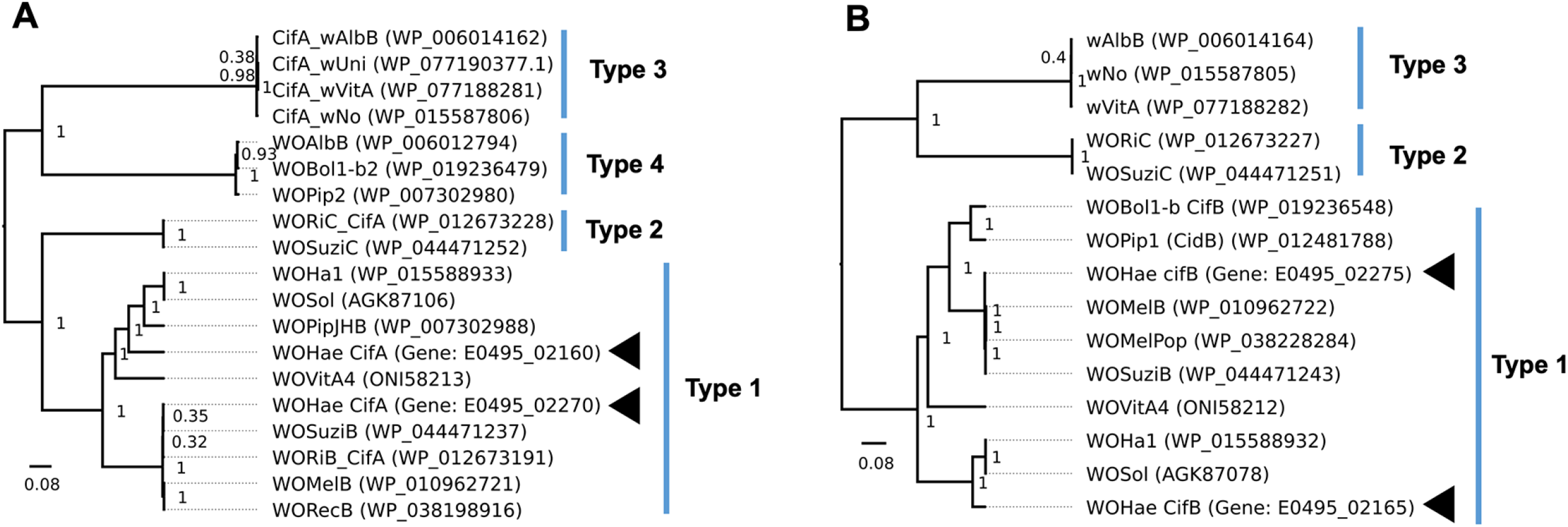
The *Wolbachia* endosymbiont *w*Hae has horizontally acquired a second cytoplasmic incompatibility loci. Maximum clade credibility (MCC) tree resulting from BEAST analyses of A) *cifA* and B) *cifB* homologues with Type numbers as designated by Lindsey et al. (2018). Posterior probability values are indicated at the nodes. *w*Hae CI genes indicated by arrowheads and branch lengths (genetic distances).

The CI genes of *Wolbachia* have been grouped into four different phylogenetic groups (Type I - IV) (19, 79), as such, we conducted a phylogenetic analysis of the complete CI genes of *w*Hae (Fig. 4). For one set of CI genes located within the WO region of *w*Hae, both copies of *cifA* and *cifB* genes were closely related to type I CifA/B proteins and closely related to *w*Ri and *w*Mel. However, it seems they have been horizontally acquired in the *w*Hae genome from other distantly related *Wolbachia*, although they cluster in Type I phylogenetic group. This report is similar to another independent acquisition of CI genes in the *Wolbachia* endosymbionts of the *Drosophila yakuba* clade which cause weak intra-and interspecific CI (80). Cooper et al. (2019) assembled the genomes of *w*Yak variants and demonstrated that while there appears to be another CI locus in these genomes, the presence of an inversion introduces several stop codons within the cidB^wYak-clade^ locus relative to the same region in cidB^wMel^, speculated to potentially render this gene non-functional (81). By comparison, both genes within the CI loci in *w*Hae are seemingly complete with no premature stops and presumed to encode for functional proteins. Previous studies have suggested that the CI gene sets *cifA* and *cifB* vary in copy number across CI-inducing *Wolbachia* strains and are directly correlated with the extent of CI (strong or weak) (79). The acquisition of a second set of CI genes corroborates previously unpublished experiments conducted where *w*Hae *Wolbachia* from the Kerrville reference strain demonstrated a strong CI phenotype (personal communication Felix Guerrero; USDA-lab, US). The transcriptional activity of the CI genes have previously been explored by Lindsey et al. (2018) who demonstrated that both *cifA* and *cifB* show differential transcriptional activity across host development (82). Again, RNA-Seq data of all life stages were mapped to the *w*Hae genome and we examined the mapped reads at these two CI loci. Reads mapped exclusively to one CI region and very few reads mapped to both (MAPQ score 0). In general, the *cifA* gene was more transcriptionally active than the *cifB* gene in both loci, as also previously reported (Fig. 4) (79). The evidence of two transcriptionally active CI loci may explain the high incidence of *Wolbachia* in wild-caught specimens of *H. i. irritans* as *Wolbachia* has been identified in 100% of all collected individuals from Hungry (10/10) (32), as well as all 15 tested horn flies from two wild locations in Alberta, Canada and also in 54/55 individuals tested in two independent screens of the laboratory colony of Lethbridge Research Centre, Alberta, Canada (27).

### The Kerrickville *Wolbachia w*Hae strain is closely related to wild *H. i. irritans Wolbachia* strains from the US, Mexico, Canada and Hungary

Previous publications have demonstrated the presence of *Wolbachia* from wild-caught and laboratory colonies of *H. i. irritans* through amplicon sanger sequencing of samples (30, 32) (83) or identifying *Wolbachia* reads in pyrosequencing-based approaches or expressed sequence tags (EST) (28, 29). A survey of currently deposited Genbank data of sequenced amplicons from *Wolbachia* endosymbionts of *H. i. irritans* are limited to partial fragments of the *Wolbachia surface protein* (*wsp*) gene (30, 83) or fragments of the *16S ribosomal RNA* gene (32). BLASTn analysis of the *wsp* fragment sample of the Kerrickville colony used by Jeyaprakash and Hoy (2000), designated as wIrr-A1 (Genbank: AF217714.1) (30), showed 100% identity with the *wsp* locus of *w*Hae (Gene: E0495; Position: 1,282,799-1,283,488) over a 548bp region. Similar high nucleotide identity of the wsp fragment of *H. i. irritans* samples, originating from Lethbridge, Alberta, Canada designated *w*Irr (Genbank: DQ380856.1), with the *w*Hae *wsp* was found; 99.64% with only two nucleotide differences over an amplicon of 554bp. In addition, the *w*Hae *16s rRNA* gene (Position: 882,502-884,006) and partial 16S rRNA fragments from two *Wolbachia* strains from *H. i. irritans* Hungary samples (Isolate G25 Genbank: EU315781.1, Isolate G24 Genbank: EU315780.1) were 99.62% identical with 264/265 sequence similarity. While this suggests the *w*Hae strain of *Wolbachia* is very closely related to the Canadian and Hungarian *H. i. irritans* samples, the nature of the amplicon size and the high nucleotide identity between strains makes it difficult to state this with complete certainty.

As high-throughput sequencing allows for a closer examination of relatedness between the Kerrickville *w*Hae *Wolbachia* strain and wild-caught *H. i. irritans* harbouring *Wolbachia*, we re-analysed EST, DNA-Seq and RNA-Seq data from a number of publications using wild-caught flies from Mexico, USA and also Uruguay (Table 4). Using our assembled genome as a reference, we conducted BLASTn of assembled EST fragments and RNA-Seq data. We identified five EST fragments, and 394 assembled *Wolbachia* RNA contigs from wild-caught *H. i. irritans* from two different studies of Louisiana State University Agricultural Center St. Gabriel Research Station (LA, USA) (84, 85), and four EST fragments from a cattle farm in Ciudad Victoria, Tamaulipas, Mexico (86). Additionally, in six RNA-Seq libraries of newly emerged male and female horn flies wild-caught in Louisiana, USA on average 10% of each library could be mapped to the *w*Hae genome (Table S1) (87) suggesting high transcriptional activity in wild populations. All identified contigs shared closer nucleotide identity to the *w*Hae strain than any other *Wolbachia* genome deposited on NCBI (data not shown). Interestingly, we could not identify any assembled contigs or reads that mapped to the *w*Hae genome from salivary gland and midgut samples originating from wild-collected *H. i. irritans* from Canelones, Uruguay (88) suggesting that either *Wolbachia* is present in very low abundance in these samples or completely absent. In examination of the 454 DNA-Seq data originating from a single male *H. i. irritans* collected in 2003 from the Pressler Cattle Ranch in Kerrville, Texas, USA (89), of the 394,263 reads in the library, 4,581 reads could be mapped to the *w*Hae genome representing ~1.16% of the library. Subsequent *de novo* assembly yielded 1,130 assembled contigs and of which 74 were identified through BLASTn analysis as having closest bit score hit to the *w*Hae genome. The Kerrville reference *H. i. irritans* strain is a closed fly colony, which has been maintained at the USDA-ARS Knipling-Bushland U.S. Livestock Insects Research Laboratory since 1961 (33). As very few *Wolbachia* genome fragments were conserved from assembled RNA-Seq and DNA-Seq wild-caught samples, we could not construct a single phylogenetic tree for all the samples. However, the close identity of all available transcriptome and genomic data of wild-caught *H. i. irritans* flies from North American populations including Mexican to the *Wolbachia* Kerrville reference *H. i. irritans* strain suggest that likely they are also infected with the same *w*Hae strain.

**Table 4:** Metadata of available sequencing data of *H. i. irritans* samples

| Location of *H. i. irritans* sample & collection date | Sample & type of sequencing | NCBI accession | Reference |
| --- | --- | --- | --- |
| Agricultural Center St. Gabriel Research Station Louisiana, USA<br>Collected: 2008 | Larval and embryonic samples,<br>PolyA enriched RNA-Seq<br>EST | Larval EST:<br>FD457983-FD466257<br>Embryonic EST:<br>FD449556-FD457982 | (85) |
| Agricultural Center St. Gabriel Research Station Louisiana, USA<br>Collected: 28 July 2010 | Whole male and females.<br>Permethrin treated surviving males and Permethrin + Piperonyl Butoxide treated killed males<br>PolyA enriched RNA-Seq<br>Illumina Genome Analyzer II/<br>Illumina HiSeq 2000 | Assembled transcriptome accession:<br>GGLM01000000<br>Bioproject accession:<br>PRJNA429442 | (84) |
| Agricultural Center St. Gabriel Research Station Louisiana, USA<br>Collected: 2010 | Eggs, larvae, whole male and females.<br>PolyA enriched RNA-Seq<br>454 | Male: SRR003192<br>Female: SRR003191<br>Egg: SRR003190<br>Larvae: SRR003189 | (97) |
| Ciudad Victoria, Tamaulipas, Mexico<br>Collected: prior to August 2010 | Abdominal tissues of partially fed adult female,<br>PolyA enriched RNA-Seq<br>EST | HO000420-HO001165<br>HO004499-HO004744 | (86) |
| Pressler Cattle Ranch Kerrville, Texas, USA<br>Originally collected: 2003<br>Sampled: 2010 | Single male adult,<br>Random DNA sequenced using<br>454 | SRA: SRR1578740 | (89) |
| Canelones, Uruguay<br>Collected: 2016 | Salivary glands and midgut samples,<br>PolyA enriched RNA-Seq<br>Illumina HiSeq 2000 | SRA Salivary glands:<br>SRR5136552, SRR5136553<br>SRA midguts: SRR5136554,<br>SRR5136555 | (88) |

**Figure 6:**
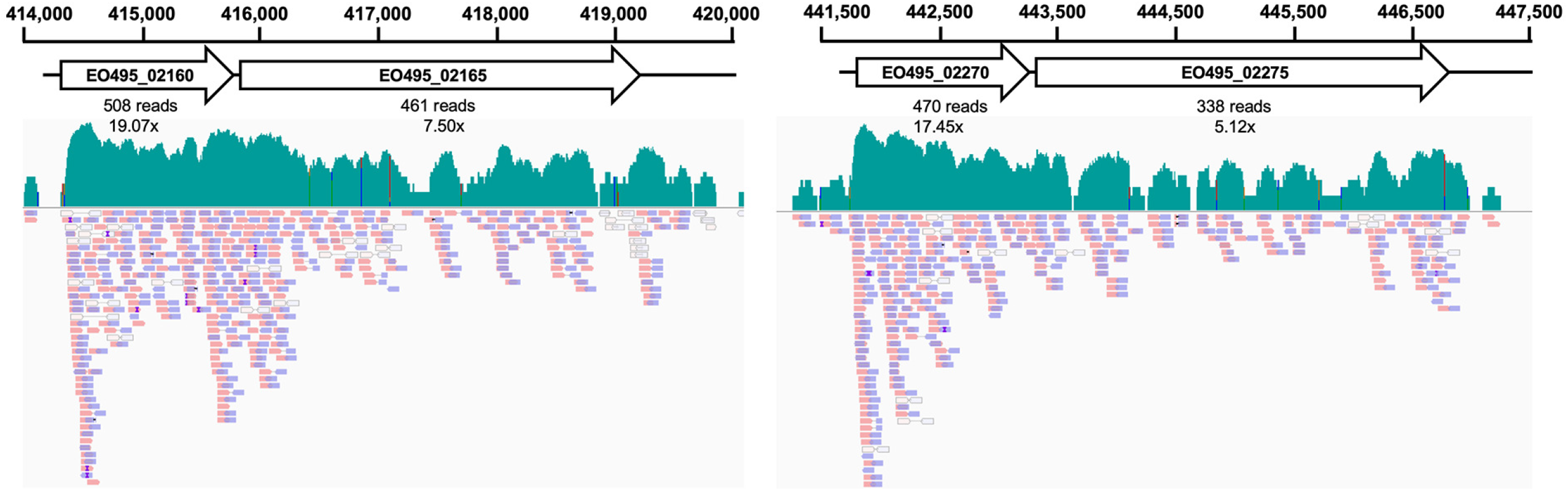
Both cytoplasmic incompatibility loci are transcriptionally active in the *Wolbachia* strain of *w*Hae. Pooled RNA-Seq reads originating from all tissues and developmental stages of all *H. i. irritans* libraries were mapped to identified CI loci in *w*Hae genomes. Resultant BAM files were visualized with Integrated Genomics Viewer (IGV v 2.5.2). Forward mapped reads are shown in red, reverse orientation reads are shown in blue. Light blue and red reads indicate a mapping quality number of 0 (MAPQ=0) which indicates that the read maps to multiple regions on the genome.

## Conclusion

In this study, we assembled and annotated a high-quality genome of the *Wolbachia* endosymbiont of *H. i. irritans* designated *w*Hae. Phylogenetic analysis of the *w*Hae strain suggests that the *w*Hae belongs to a well-supported supergroup A lineage that includes the well-studied *w*Mel, *w*Au, and *w*Ri *Wolbachia* strains from *Drosophila* spp. Comparative genomics of *w*Hae indicated acquisition of additional transcriptionally active CI loci. Phylogenetic analysis indicates either horizontal acquisition of these genes from a closely related *Wolbachia* strain or the potential loss of CI loci in other *Wolbachia* strains infecting *Drosophila spp*. The *w*Hae genome has undergone significant reassortment compared to closely related and completely assembled strains. Additional analysis of available and deposited sequencing data from wild-caught and laboratory *H. i. irritans* colonies suggest that *w*Hae is the most closely related to wild USA and Mexican samples and close relative of Canadian and Hungarian samples. This study provides the foundation for future functional studies of effects *Wolbachia* may have on life-history traits of *H. i. irritans* such as insecticide resistance and evaluating contribution of *w*Hae towards population control.

## Materials and Methods

### Genomic DNA and RNA Sequencing data

The Kerrville reference *H. i. irritans* strain is a closed fly colony which has been maintained at the USDA-ARS Knipling-Bushland U. S. Livestock Insects Research Laboratory since 1961 (33). Genomic DNA from unfed adult flies of mixed-sex originating from this strain was subjected to whole-genome sequencing, and previously deposited on the National Center for Biotechnology Information Short Read Archive (SRA) (Accession number: PRJNA30967) (33). Briefly, this data includes two PacBio runs; one 10 kb and two 20 kb insert libraries. 10 kb libraries were sequenced using C2 chemistry and P4 polymerase, whereas C3 chemistry, and P5 polymerase were used for both 20kb libraries with 3 hours of movie time. 10 kb libraries and two of the 20 kb libraries were sequenced on 12 SMRTCells, four SMRTCells, and eight SMRTCells, respectively, and all the sequences were finally pooled and uploaded under the same accession (SRA: SRR6231657). For Illumina sequencing, one short-insert paired-end library and one mate-paired end library with 6-12 kb insert size were sequenced as 100nt paired ends on the HiSeq2000 and uploaded under the same accession number (SRA: SRR6231656). Additional RNA sequencing data from different life stages and tissues of the horn flies were sequenced on a Illumina HiSeq 2000 using 2x 100nt configuration and available with the above Illumina read accession number (SRA: SRR6231656).

### *Wolbachia* genome assembly and polishing

Raw fastq files originating from Illumina and PacBio sequencing data were imported to the Galaxy Australia webserver (https://usegalaxy.org.au/). The Nextra universal transpose Illumina sequence adapters were removed and reads were quality trimmed using Trimmomatic (Galaxy version: 0.36.4) under the following conditions (Sliding window=4, average quality=20) (90). Resultant clean reads were mapped to the genome of *w*Ri (34), and *w*Au (35) using BWA-MEM (Galaxy Version 0.7.17.1) (91) under default parameters and under simple Illumina mode and PacBio mode (-x pacbio) for subsequent libraries. Mapped reads were extracted using a BAM filter (Galaxy Version 0.5.9) and were then assembled using Unicycler (Galaxy Version 0.4.1.1) (40).

### Genome annotation and comparative genomics

Coding regions and ncRNAs of the assembled *w*Hae genome contig were annotated using the NCBI prokaryotic genome annotation pipeline (42). To assess the quality of the assembly, BUSCO v. 3.1.0 was used to search for orthologs of the near-universal, single-copy genes in the BUSCO proteobacteria database (43). As a control, we performed the same search using the reference genomes for *w*Ri (34), *w*Au (35), *w*Mel (38), *w*Ha, and *w*No (37) as well as the complete *w*AlbB genome (36). Identification of phage and prophage regions of *w*Hae was conducted using the PHASTER web platform (https://phaster.ca/) (73). Groupings of orthologous clusters were identified using the Orthovenn2 web server (https://orthovenn2.bioinfotoolkits.net/) (45) under the following conditions: E-value: 1e-2, Inflation value: 1.5. Insertion sequence (IS) elements of *w*Hae were identified using the ISsaga web server platform (http://issaga.biotoul.fr/) (92). For nucleotide synteny plots of *w*Hae MAFFT (https://mafft.cbrc.jp/alignment/server/) (93) was used to align *w*Hae and other genomes and then visualised by dot-plots of matches (without extensions) identified using the LAST algorithm which compares sequences by adaptive and fixed-length seeds (score=39, E=8.4e-11). Comparisons between the putative prophage regions of *w*Hae were examined using BLASTn and visualised using Easyfig (94).

### Phylogenetic analyses of *w*Hae and cytoplasmic incompatibility loci

For full genome phylogenetic analyses, we used 79 non-recombinant gene loci, which has been previously determined by Bleidorn & Gerth (2018) to perform well from 19 strains of *Wolbachia* (51). These were downloaded (https://github.com/gerthmicha/wolbachia-mlst), aligned using MUSCLE (95) and concatenated. The resultant alignment was analysed using Bayesian evolutionary analysis by sampling trees (BEAST v2.5.1) (96), split into individual codon positions with linked site model, and unlinked clock model under the General Time Reversible and Gamma = 4 nucleotide substitution model. Clock rates were drawn from a log-normal distribution. Additional parameters were a chain length of 10 million steps sampling every 10,000 steps under a Yule model. For phylogenetic placement of the CI genes within *w*Hae, identified Cif homologs were first aligned using MUltiple Sequence Comparison by Log-Expectation (MUSCLE) (95) and also subjected to BEAST (96) with 10 million steps with a pre-burnin of 100,000 with sampling being conducted every 20,000 steps under a Yule model and a general empirical model of protein evolution (WAG) amino acid substitution model. For both BEAST runs convergence for all parameters as well as stationary distributions of the MCMC chain were inspected using Tracer v1.7.1 (effective sample sizes of >400). The maximum clade credibility (MCC) tree (i.e. the tree with the largest product of posterior clade probabilities) was selected from the posterior tree distribution using the program TreeAnnotator (included in the BEAST package) after a 10% burn in. Resultant MCC trees were then visualised using FigTree v1.4.4.

### Data availability and accession numbers

PacBio and Illumina raw sequencing data are available from the NCBI short read archive under accession numbers SRR6231657 and SRR6231656, respectively. The assembled *Wolbachia pipientis w*Hae strain has been deposited in Genbank under the accession number CP037426. Additional sequencing data and metadata used for validation are available in Supplementary Files. Alignment files used to make the Phylogeny are available at the Figshare collection (<u>deposited in Figshare collection, will make public when submitted</u>)

## Acknowledgements

The authors acknowledge the support of Felix Guerrero from the USDA-ARS Knipling-Bushland US Livestock Insects Research Laboratory. Analysis was conducted using the Australian Galaxy platform (https://usegalaxy.org.au/) with the support and technical assistance of Igor Makunin. This project was funded by the Australian Research Council grant (DP150101782) to SA and the University of Queensland scholarship to M.M. and R.P.

---

**Figure S1: Similar BUSCO scores across all the complete *Wolbachia* genomes.** The BUSCO pipeline was used to measure the proportion of highly conserved, single copy orthologs (BUSCO groups). The set of reference BUSCO group was set to the lineage “Proteobacteria”, which contains 221 BUSCO derived from 1520 proteobacterial species.

**Figure S2: Orthologous clusters of the proteome of supergroup A *Wolbachia* strains.** Analysis conducted using Orthovenn webserver under the conditions: E-value: 1e-2, Inflation value: 15.

## References

1. Changbunjong T, Weluwanarak T, Samung Y, Ruangsittichai J. 2016. Molecular identification and genetic variation of hematophagous flies, (Diptera: Muscidae: Stomoxyinae) in Thailand based on cox1 barcodes. J Asia-Pac Entomol 19:1117–1123.

2. Oyarzún M, Quiroz A, Birkett M. 2008. Insecticide resistance in the horn fly: alternative control strategies. Med Vet Entomol 22:188–202.

3. Cupp E, Cupp M, Ribeiro J, Kunz S. 1998. Blood-feeding strategy of *Haematobia irritans* (Diptera: Muscidae). J Med Entomol 35:591–595.

4. Grisi L, Leite RC, Martins JR, Barros AT, Andreotti R, Cancado PH, Leon AA, Pereira JB, Villela HS. 2014. Reassessment of the potential economic impact of cattle parasites in Brazil. Rev Bras Parasitol Vet 23:150–156.

5. Oyarzún M, Quiroz A, Birkett M. 2008. Insecticide resistance in the horn fly: alternative control strategies. Med Vet Entomol 22:188–202.

6. Lane J, Jubb T, Shephard R, Webb-Ware J, Fordyce G. 2015. Priority list of endemic diseases for the red meat industries. Meat & Livestock Australia (report) B.AHE.0010.

7. Schnitzerling H, Noble P, Macqueen A, Dunham R. 1982. Resistance of the buffalo fly, *Haematobia irritans exigua* (De Meijere), to two synthetic pyrethroids and DDT. Aust J Entomol 21:77–80.

8. Rothwell J, Morgan J, James P, Brown G, Guerrero F, Jorgensen W. 2011. Mechanism of resistance to synthetic pyrethroids in buffalo flies in south-east Queensland. Aust Vet J 89:70–72.

9. Zug R, Hammerstein P. 2012. Still a host of hosts for *Wolbachia*: analysis of recent data suggests that 40% of terrestrial arthropod species are infected. PLoS One 7:e38544.

10. LePage D, Bordenstein SR. 2013. *Wolbachia*: can we save lives with a great pandemic? Trends Parasitol 29:385–393.

11. Caragata EP, Dutra HL, Moreira LA. 2016. Exploiting intimate relationships: controlling mosquito-transmitted disease with *Wolbachia*. Trends Parasitol 32:207–218.

12. Beckmann JF, Bonneau M, Chen H, Hochstrasser M, Poinsot D, Merçot H, Weill M, Sicard M, Charlat S. 2019. The toxin-antidote model of cytoplasmic incompatibility: genetics and evolutionary implications. Trends Genet 35: 175–185.

13. Werren JH, Baldo L, Clark ME. 2008. *Wolbachia*: master manipulators of invertebrate biology. Nat Rev Micriobiol 6:741.

14. Poinsot D, Charlat S, Mercot H. 2003. On the mechanism of *Wolbachia*-induced cytoplasmic incompatibility: Confronting the models with the facts. Bioessays 25:259–265.

15. Landmann F, Orsi GA, Loppin B, Sullivan W. 2009. *Wolbachia*-mediated cytoplasmic incompatibility is associated with impaired histone deposition in the male pronucleus. PLoS Pathogens 5:e1000343.

16. Tram U, Sullivan W. 2002. Role of delayed nuclear envelope breakdown and mitosis in *Wolbachia*-induced cytoplasmic incompatibility. Science 296:1124–1126.

17. Tram U, Fredrick K, Werren JH, Sullivan W. 2006. Paternal chromosome segregation during the first mitotic division determines *Wolbachia*-induced cytoplasmic incompatibility phenotype. J Cell Sci 119:3655–3663.

18. Callaini G, Dallai R, Riparbelli MG. 1997. Wolbachia-induced delay of paternal chromatin condensation does not prevent maternal chromosomes from entering anaphase in incompatible crosses of Drosophila simulans. J Cell Sci 110:271–280.

19. LePage DP, Metcalf JA, Bordenstein SR, On J, Perlmutter JI, Shropshire JD, Layton EM, Funkhouser-Jones LJ, Beckmann JF, Bordenstein SR. 2017. Prophage genes recapitulate and enhance *Wolbachia*-induced cytoplasmic incompatibility. Nature 543:243.

20. Beckmann JF, Ronau JA, Hochstrasser M. 2017. A Wolbachia deubiquitylating enzyme induces cytoplasmic incompatibility. Nat Microbiol 2:17007.

21. Shropshire JD, On J, Layton EM, Zhou H, Bordenstein SR. 2018. One prophage WO gene rescues cytoplasmic incompatibility in Drosophila melanogaster. Proc Natl Acad Sci, USA 115:4987–4991.

22. Jeffries CL, Walker T. 2016. *Wolbachia* biocontrol strategies for arboviral diseases and the potential influence of resident *Wolbachia* strains in mosquitoes. Curr Trop Med Rep 3:20–25.

23. Iturbe-Ormaetxe I, Walker T, O’Neill SL. 2011. Wolbachia and the biological control of mosquito-borne disease. EMBO Rep 12:508–518.

24. Berticat C, Rousset F, Raymond M, Berthomieu A, Weill M. 2002. High *Wolbachia* density in insecticide-resistant mosquitoes. Proc Biol Sci 269:1413–6.

25. Duron O, Labbe P, Berticat C, Rousset F, Guillot S, Raymond M, Weill M. 2006. High *Wolbachia* density correlates with cost of infection for insecticide resistant *Culex pipiens* mosquitoes. Evolution 60:303–14.

26. Echaubard P, Duron O, Agnew P, Sidobre C, Noel V, Weill M, Michalakis Y. 2010. Rapid evolution of *Wolbachia* density in insecticide resistant *Culex pipiens*. Heredity 104:15–19.

27. Zhang B, McGraw E, Floate KD, James P, Jorgensen W, Rothwell J. 2009. *Wolbachia* infection in Australasian and North American populations of *Haematobia irritans* (Diptera: Muscidae). Vet Parasitol 162:350–353.

28. Torres L, Almazán C, Ayllón N, Galindo RC, Rosario-Cruz R, Quiroz-Romero H, Gortazar C, de la Fuente J. 2012. Identification of microorganisms in partially fed female horn flies, *Haematobia irritans*. Parasitol Res 111:1391–1395.

29. Palavesam A, Guerrero FD, Heekin AM, Wang J, Dowd SE, Sun Y, Foil LD, de León AAP. 2012. Pyrosequencing-based analysis of the microbiome associated with the horn fly, *Haematobia irritans*. PLoS One 7:e44390.

30. Jeyaprakash A, Hoy MA. 2000. Long PCR improves *Wolbachia* DNA amplification: wsp sequences found in 76% of sixty-three arthropod species. Insect Mol Biol 9:393–405.

31. Floate KD, Kyei-Poku GK, Coghlin PC. 2006. Overview and relevance of Wolbachia bacteria in biocontrol research. Biocontrol Sci Technol16:767–788.

32. Hornok S, Földvári G, Elek V, Naranjo V, Farkas R, de la Fuente J. 2008. Molecular identification of *Anaplasma marginale* and rickettsial endosymbionts in blood-sucking flies (Diptera: Tabanidae, Muscidae) and hard ticks (Acari: Ixodidae). Vet Parasitol 154:354–359.

33. Konganti K, Guerrero FD, Schilkey F, Ngam P, Jacobi JL, Umale PE, de Leon AAP, Threadgill DW. 2018. A whole genome assembly of the Horn fly, *Haematobia irritans*, and prediction of genes with roles in metabolism and sex determination. G3: Genes, Genomes, Genetics 8:1675–1686.

34. Klasson L, Westberg J, Sapountzis P, Näslund K, Lutnaes Y, Darby AC, Veneti Z, Chen L, Braig HR, Garrett R. 2009. The mosaic genome structure of the *Wolbachia w*Ri strain infecting *Drosophila simulans*. Proc Natl Acad Sci, USA 106:5725–5730.

35. Sutton ER, Harris SR, Parkhill J, Sinkins SP. 2014. Comparative genome analysis of *Wolbachia* strain *w*Au. BMC Genomics 15:928.

36. Sinha A, Li Z, Sun L, Carlow CK. 2019. Complete genome sequence of the *Wolbachia w*AlbB endosymbiont of *Aedes albopictus*. Genome Biol Evo 11:706–720.

37. Ellegaard KM, Klasson L, Näslund K, Bourtzis K, Andersson SG. 2013. Comparative genomics of *Wolbachia* and the bacterial species concept. PLoS Genetics 9:e1003381.

38. Wu M, Sun LV, Vamathevan J, Riegler M, Deboy R, Brownlie JC, McGraw EA, Martin W, Esser C, Ahmadinejad N. 2004. Phylogenomics of the reproductive parasite *Wolbachia pipientis w*Mel: a streamlined genome overrun by mobile genetic elements. PLoS Biol 2:e69.

39. Li H, Durbin R. 2010. Fast and accurate long-read alignment with Burrows-Wheeler transform. Bioinformatics 26:589–95.

40. Wick RR, Judd LM, Gorrie CL, Holt KE. 2017. Unicycler: resolving bacterial genome assemblies from short and long sequencing reads. PLoS Comput Biol 13:e1005595.

41. Walker BJ, Abeel T, Shea T, Priest M, Abouelliel A, Sakthikumar S, Cuomo CA, Zeng Q, Wortman J, Young SK. 2014. Pilon: an integrated tool for comprehensive microbial variant detection and genome assembly improvement. PLoS One 9:e112963.

42. Tatusova T, DiCuccio M, Badretdin A, Chetvernin V, Nawrocki EP, Zaslavsky L, Lomsadze A, Pruitt KD, Borodovsky M, Ostell J. 2016. NCBI prokaryotic genome annotation pipeline. Nucleic Acids Res 44:6614–6624.

43. Waterhouse RM, Seppey M, Simão FA, Manni M, Ioannidis P, Klioutchnikov G, Kriventseva EV, Zdobnov EM. 2017. BUSCO applications from quality assessments to gene prediction and phylogenomics. Mol Biol Evol 35:543–548.

44. Seppey M, Manni M, Zdobnov EM. 2019. BUSCO: Assessing genome assembly and annotation completeness. Methods Mol Biol 1962:227–245.

45. Wang Y, Coleman-Derr D, Chen G, Gu YQ. 2015. OrthoVenn: a web server for genome wide comparison and annotation of orthologous clusters across multiple species. Nucleic Acids Res 43:W78–W84.

46. Hertig M. 1936. The rickettsia, *Wolbachia pipientis* (gen. et sp. n.) and associated inclusions of the mosquito, *Culex pipiens*. Parasitology 28:453–486.

47. Lo N, Paraskevopoulos C, Bourtzis K, O’neill S, Werren J, Bordenstein S, Bandi C. 2007. Taxonomic status of the intracellular bacterium *Wolbachia pipientis*. Int J Syst Evol Microbiol 57:654–657.

48. Lindsey AR, Bordenstein SR, Newton IL, Rasgon JL. 2016. *Wolbachia pipientis* should not be split into multiple species: A response to Ramírez-Puebla et al.,“Species in Wolbachia? Proposal for the designation of ‘Candidatus Wolbachia bourtzisii’,‘Candidatus Wolbachia onchocercicola’,‘Candidatus Wolbachia blaxteri’,‘Candidatus Wolbachia brugii’,‘Candidatus Wolbachia taylori’,‘Candidatus Wolbachia collembolicola’ and ‘Candidatus Wolbachia multihospitum’for the different species within *Wolbachia* supergroups”. Syst Appl Microbiol 39:220.

49. Ramírez-Puebla ST, Servín-Garcidueñas LE, Ormeño-Orrillo E, de León AV-P, Rosenblueth M, Delaye L, Martínez J, Martínez-Romero E. 2015. Species in Wolbachia? Proposal for the designation of ‘Candidatus Wolbachia bourtzisii’,‘Candidatus Wolbachia onchocercicola’,‘Candidatus Wolbachia blaxteri’,‘Candidatus Wolbachia brugii’,‘Candidatus Wolbachia taylori’,‘Candidatus Wolbachia collembolicola’and ‘Candidatus Wolbachia multihospitum’for the different species within *Wolbachia* supergroups. Syst Appl Microbiol 38:390–399.

50. Baldo L, Dunning Hotopp JC, Jolley KA, Bordenstein SR, Biber SA, Choudhury RR, Hayashi C, Maiden MC, Tettelin H, Werren JH. 2006. Multilocus sequence typing system for the endosymbiont *Wolbachia pipientis*. Appl Environ Microbiol 72:7098–110.

51. Bleidorn C, Gerth M. 2017. A critical re-evaluation of multilocus sequence typing (MLST) efforts in Wolbachia. FEMS Microbiol Ecol 94:fix163.

52. Turelli M, Cooper BS, Richardson KM, Ginsberg PS, Peckenpaugh B, Antelope CX, Kim KJ, May MR, Abrieux A, Wilson DA. 2018. Rapid global spread of *w*Ri-like *Wolbachia* across multiple *Drosophila*. Curr Biol 28:963–971. e8.

53. Pietri JE, DeBruhl H, Sullivan W. 2016. The rich somatic life of *Wolbachia*. Microbiology Open 5:923–936.

54. Oliveira MT, Barau JG, Junqueira ACM, Feijão PC, da Rosa AC, Abreu CF, Azeredo-Espin AML, Lessinger AC. 2008. Structure and evolution of the mitochondrial genomes of Haematobia irritans and *Stomoxys calcitrans*: the Muscidae (Diptera: Calyptratae) perspective. Mol Phylogenet Evol 48:850–857.

55. Ding S, Li X, Wang N, Cameron SL, Mao M, Wang Y, Xi Y, Yang D. 2015. The phylogeny and evolutionary timescale of muscoidea (Diptera: Brachycera: Calyptratae) inferred from mitochondrial genomes. PLoS One 10:e0134170.

56. Bandi C, Anderson TJ, Genchi C, Blaxter ML. 1998. Phylogeny of *Wolbachia* in filarial nematodes. Proc Biol Sci 265:2407–13.

57. Werren JH, Zhang W, Guo LR. 1995. Evolution and phylogeny of *Wolbachia*: reproductive parasites of arthropods. Proc Biol Sci 261:55–63.

58. Gerth M, Bleidorn C. 2016. Comparative genomics provides a timeframe for *Wolbachia* evolution and exposes a recent biotin synthesis operon transfer. Nat Microbiol 2:16241.

59. Gillings MR. 2017. Lateral gene transfer, bacterial genome evolution, and the Anthropocene. Ann NY Acad Sci 1389:20–36.

60. Kent BN, Salichos L, Gibbons JG, Rokas A, Newton IL, Clark ME, Bordenstein SR. 2011. Complete bacteriophage transfer in a bacterial endosymbiont (*Wolbachia*) determined by targeted genome capture. Genome Biol Evol 3:209–218.

61. Frost LS, Leplae R, Summers AO, Toussaint A. 2005. Mobile genetic elements: the agents of open source evolution. Nat Rev Microbiol 3:722.

62. Newton IL, Bordenstein SR. 2011. Correlations between bacterial ecology and mobile DNA. Curr Microbiol 62:198–208.

63. Rozewicki J, Li S, Amada KM, Standley DM, Katoh K. 2019. MAFFT-DASH: integrated protein sequence and structural alignment. Nucleic Acids Res 47: W5–W10.

64. Metcalf JA, Jo M, Bordenstein SR, Jaenike J, Bordenstein SR. 2014. Recent genome reduction of *Wolbachia* in *Drosophila recens* targets phage WO and narrows candidates for reproductive parasitism. PeerJ 2:e529.

65. Klasson L, Walker T, Sebaihia M, Sanders MJ, Quail MA, Lord A, Sanders S, Earl J, O’neill SL, Thomson N. 2008. Genome evolution of *Wolbachia* strain *w*Pip from the *Culex pipiens* group. Mol Biol Evol 25:1877–1887.

66. Foster J, Ganatra M, Kamal I, Ware J, Makarova K, Ivanova N, Bhattacharyya A, Kapatral V, Kumar S, Posfai J. 2005. The *Wolbachia* genome of *Brugia malayi*: endosymbiont evolution within a human pathogenic nematode. PLoS Biol 3:e121.

67. Newton IL, Clark ME, Kent BN, Bordenstein SR, Qu J, Richards S, Kelkar YD, Werren JH. 2016. Comparative genomics of two closely related Wolbachia with different reproductive effects on hosts. Genome Biol Evol 8:1526–1542.

68. Siguier P, Pérochon J, Lestrade L, Mahillon J, Chandler M. 2006. ISfinder: the reference centre for bacterial insertion sequences. Nucleic Acids Res 34:D32–D36.

69. Hibler CP. 1966. Development of *Stephanofilaria stilesi* in the horn fly. J Parasitol 52:890–898.

70. Kent BN, Bordenstein SR. 2010. Phage WO of *Wolbachia*: lambda of the endosymbiont world. Trends Microbiol 18:173–181.

71. Bordenstein SR, Bordenstein SR. 2016. Eukaryotic association module in phage WO genomes from *Wolbachia*. Nat Commun 7:13155.

72. Tanaka K, Furukawa S, Nikoh N, Sasaki T, Fukatsu T. 2009. Complete WO phage sequences reveal their dynamic evolutionary trajectories and putative functional elements required for integration into the *Wolbachia* genome. Appl Environ Microbiol 75:5676–5686.

73. Arndt D, Grant JR, Marcu A, Sajed T, Pon A, Liang Y, Wishart DS. 2016. PHASTER: a better, faster version of the PHAST phage search tool. Nucleic Acids Res 44:W16–W21.

74. Biliske JA, Batista PD, Grant CL, Harris HL. 2011. The bacteriophage WORiC is the active phage element in wRi of *Drosophila simulans* and represents a conserved class of WO phages. BMC Microbiol 11:251.

75. Iturbe-Ormaetxe I, Burke GR, Riegler M, O’Neill SL. 2005. Distribution, expression, and motif variability of ankyrin domain genes in *Wolbachia pipientis*. J Bacteriol 187:5136–5145.

76. Voronin DA, Kiseleva EV. 2007. Functional role of proteins containing ankyrin repeats. Tsitologiia 49:989–999.

77. Ishmael N, Dunning Hotopp JC, Ioannidis P, Biber S, Sakamoto J, Siozios S, Nene V, Werren J, Bourtzis K, Bordenstein SR, Tettelin H. 2009. Extensive genomic diversity of closely related *Wolbachia* strains. Microbiology 155:2211–2222.

78. Siozios S, Ioannidis P, Klasson L, Andersson SG, Braig HR, Bourtzis K. 2013. The diversity and evolution of *Wolbachia* ankyrin repeat domain genes. PLoS One 8:e55390.

79. Lindsey AR, Rice DW, Bordenstein SR, Brooks AW, Bordenstein SR, Newton IL. 2018. Evolutionary genetics of cytoplasmic incompatibility genes cifA and cifB in prophage WO of *Wolbachia*. Genome Biol Evol 10:434–451.

80. Cooper BS, Ginsberg PS, Turelli M, Matute DR. 2017. *Wolbachia* in the *Drosophila yakuba* complex: Pervasive frequency variation and weak cytoplasmic incompatibility, but no apparent effect on reproductive isolation. Genetics 205:333–351.

81. Cooper BS, Vanderpool D, Conner WR, Matute DR, Turelli M. 2019. *Wolbachia* acquisition by *Drosophila yakuba*-Clade hosts and transfer of incompatibility loci between distantly related *Wolbachia*. Genetics 212:1399–1419.

82. Lindsey AR, Rice DW, Bordenstein SR, Brooks AW, Bordenstein SR, Newton IL. 2018. Evolutionary genetics of cytoplasmic incompatibility genes cifA and cifB in prophage WO of *Wolbachia*. Genome Biol Evo 10:434–451.

83. Kyei-Poku G, Giladi M, Coghlin P, Mokady O, Zchori-Fein E, Floate K. 2006. Wolbachia in wasps parasitic on filth flies with emphasis on Spalangia cameroni. Entomol Exp Appl 121:123–135.

84. Domingues LN, Guerrero FD, Cameron C, Farmer A, Bendele KG, Foil LD. 2018. The assembled transcriptome of the adult horn fly, *Haematobia irritans*. Data in brief 19:1933–1940.

85. Guerrero F, Dowd S, Nene V, Foil L. 2008. Expressed cDNAS from embryonic and larval stages of the horn fly (Diptera: Muscidae). J Med Entomol 45:686–692.

86. Torres L, Almazán C, Ayllón N, Galindo RC, Rosario-Cruz R, Quiroz-Romero H, de la Fuente J. 2011. Functional genomics of the horn fly, *Haematobia irritans* (Linnaeus, 1758). BMC Genomics 12:105.

87. Domingues LN, Guerrero FD, Cameron C, Farmer A, Bendele KG, Foil LD. 2018. The assembled transcriptome of the adult horn fly, *Haematobia irritans*. Data Brief 19:1933–1940.

88. Ribeiro JM, Debat HJ, Boiani M, Ures X, Rocha S, Breijo M. 2019. An insight into the sialome, mialome and virome of the horn fly, *Haematobia irritans*. BMC Genomics 20:616.

89. Ramakodi MP, Singh B, Wells JD, Guerrero F, Ray DA. 2015. A 454 sequencing approach to dipteran mitochondrial genome research. Genomics 105:53–60.

90. Bolger AM, Lohse M, Usadel B. 2014. Trimmomatic: a flexible trimmer for Illumina sequence data. Bioinformatics 30:2114–2120.

91. Li H, Durbin R. 2010. Fast and accurate long-read alignment with Burrows–Wheeler transform. Bioinformatics 26:589–595.

92. Varani AM, Siguier P, Gourbeyre E, Charneau V, Chandler M. 2011. ISsaga is an ensemble of web-based methods for high throughput identification and semi-automatic annotation of insertion sequences in prokaryotic genomes. Genome Biol 12:R30.

93. Katoh K, Standley DM. 2013. MAFFT multiple sequence alignment software version 7: improvements in performance and usability. Mol Biol Evol 30:772–780.

94. Sullivan MJ, Petty NK, Beatson SA. 2011. Easyfig: a genome comparison visualizer. Bioinformatics 27:1009–10.

95. Edgar RC. 2004. MUSCLE: multiple sequence alignment with high accuracy and high throughput. Nucleic Acids Res 32:1792–1797.

96. Drummond AJ, Suchard MA, Xie D, Rambaut A. 2012. Bayesian phylogenetics with BEAUti and the BEAST 1.7. Mol Biol Evol 29:1969–1973.

97. Guerrero FD, Dowd SE, Sun Y, Saldivar L, Wiley GB, Macmil SL, Najar F, Roe BA, Foil LD. 2009. Microarray analysis of female- and larval-specific gene expression in the horn fly (Diptera: Muscidae). J Med Entomol 46:257–70.

